## Supplementary Figures for "Lamivudine (3TC), a nucleoside reverse transcriptase inhibitor, prevents the neuropathological alterations present in mutant tau transgenic mice"

**Affiliation:**
[a]Centro de Biología Molecular "Severo Ochoa", CSIC/UAM, Universidad Autónoma de Madrid, Cantoblanco, 28049 Madrid, Spain.

***Address correspondence to**: Félix Hernández, Centro de Biología Molecular "Severo Ochoa", CSIC/UAM, Universidad Autónoma de Madrid, Cantoblanco, 28049 Madrid, Spain; Tel.: +34 91 196 45 63; fax: +34 91 196 44 20.
https://orcid.org/0000-0001-8753-8249

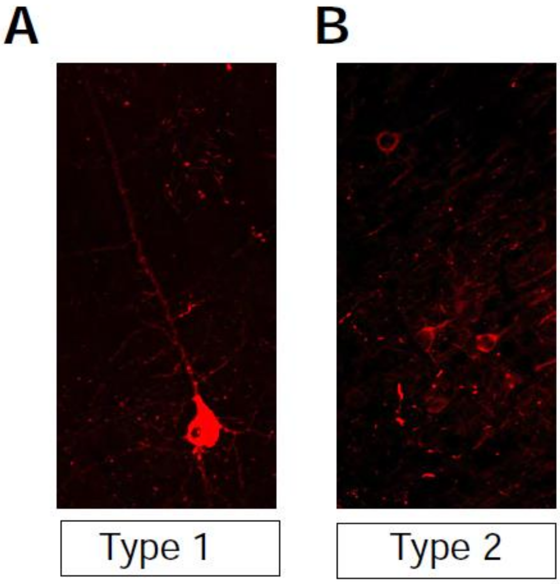


**Supplementary figure 1: Representative AT8+ Type 1 or "Tangle-like" neuron (A) and Type 2 neuron with a diffuse pattern (B)**.

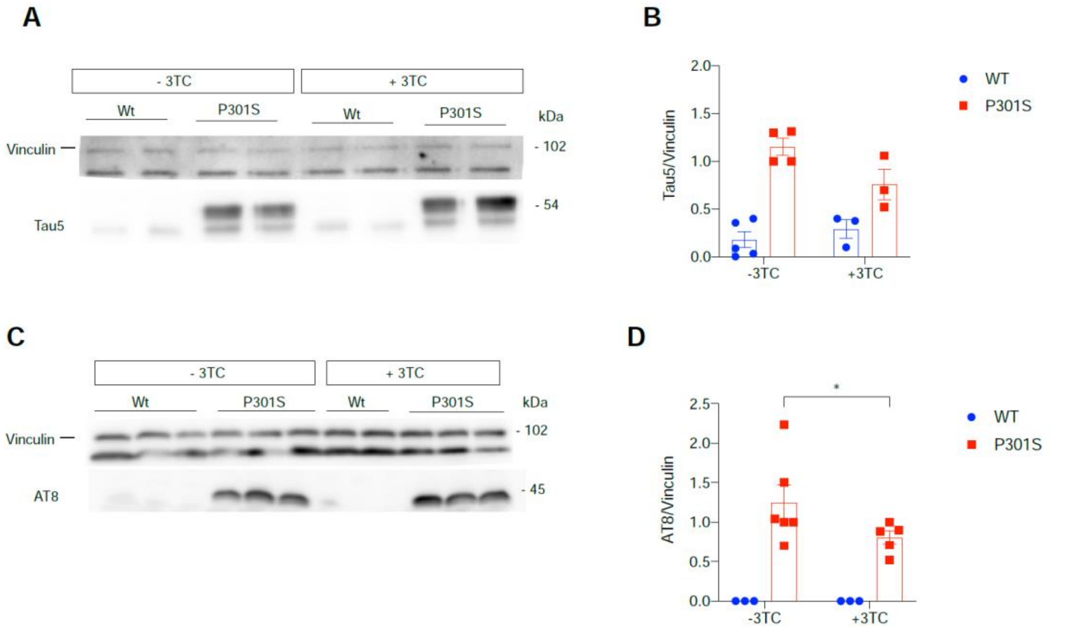


**Supplementary figure 2: Hyperphosphorylation of tau in P301S mice.** Western blot of protein extracts from the cortex of wild-type or P301S mice probed with Tau5 (A-B) and AT8 (C-D) antibodies. Vinculin blots are shown as controls of total protein loading. (B and D) Histograms show relative Tau5 and AT8 levels in untreated and 3TC-treated mice. *p<0.052 using Student's *t*-test for comparison (n=3–5).

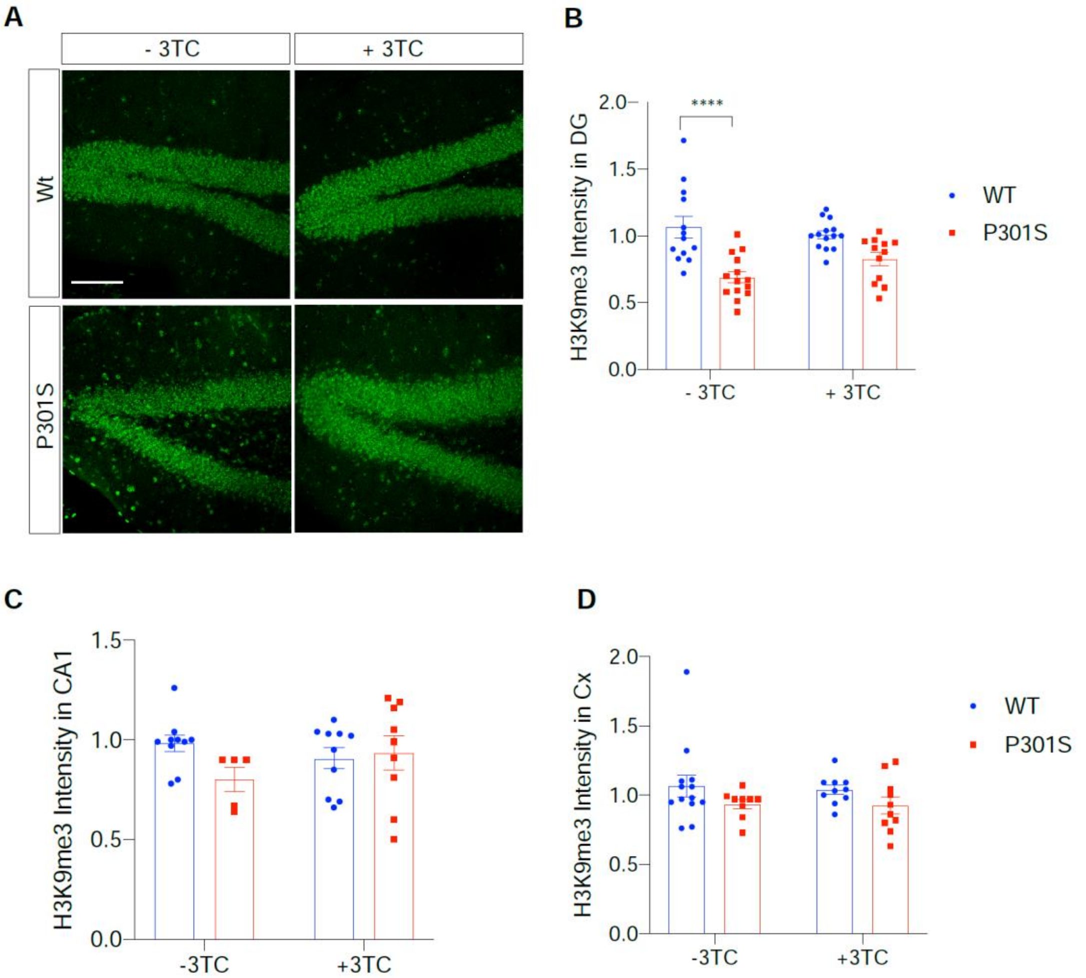


**Supplementary figure 3: 3TC treatment promotes compaction of heterochromatin in P301S mice.** (A) Representative images of the dentate gyrus with the H3K9me3 marker of slices from mice treated with 3TC and untreated counterparts and their quantification (B). The graph shows the mean± SEM (n= 13–15 per condition). ****$p<0.0001$ using two-way ANOVA followed by Student's t-test for comparisons. Scale bar, 20 µm. 3TC treatment did not promote the compaction of heterochromatin in the CA1 and cortex of wild-type or P301S mice. (C-D) Representative immunofluorescence images of CA1 and cortex with the H3K9me3 marker. Graphs show the mean± SEM (n= 13–15 per condition). Two-way ANOVA did not show statistically significant differences between the means. Scale bar, 20 µm.
